# Extensive phylogenies of human development reveal variable embryonic patterns

**DOI:** 10.1101/2020.11.25.397828

**Authors:** Tim H. H. Coorens, Luiza Moore, Philip S. Robinson, Rashesh Sanghvi, Joseph Christopher, James Hewinson, Alex Cagan, Thomas R. W. Oliver, Matthew D. C. Neville, Yvette Hooks, Ayesha Noorani, Thomas J. Mitchell, Rebecca C. Fitzgerald, Peter J. Campbell, Iñigo Martincorena, Raheleh Rahbari, Michael R. Stratton

**Affiliations:** Wellcome Sanger Institute, Hinxton, CB10 1SA, UK; Department of Pathology, University of Cambridge, Cambridge, CB2 0QQ, UK; Department of Paediatrics, University of Cambridge, Cambridge, CB2 0QQ, UK; Cambridge University Hospitals NHS Foundation Trust, Cambridge, CB2 0QQ, UK; Department of Surgery, University of Cambridge, Cambridge, CB2 0QQ, UK; MRC Cancer Unit, University of Cambridge, Biomedical Campus, Cambridge, CB2 OXZ, UK

**Author notes:** Joint first authors (Tim H. H. Coorens, Luiza Moore). Correspondence to (MRS).

## Abstract

Starting from the zygote, all cells in the developing and adult human body continuously acquire mutations. A mutation shared between two different cells implies a shared progenitor cell and can thus be used as a naturally occurring marker for lineage tracing. Here, we reconstruct extensive phylogenies of normal tissues from three adult individuals using whole-genome sequencing of 511 laser capture microdissected samples from multiple organs. Early embryonic progenitor cells inferred from the phylogenies often contribute in different proportions to the adult body and the extent of this asymmetry is variable between individuals, with ratios between the first two reconstructed cells ranging from 56:44 to 92:8. Asymmetries also pervade subsequent cell generations and can differ between tissues in the same individual. The phylogenies also resolve the spatial embryonic origins and patterning of tissues, revealing a spatial effect in the development of the human brain. Supplemented by data on eleven men, we timed the split between soma and germline, with the earliest observed segregation occurring at the first cell divisions. This research demonstrates that, despite reaching the same ultimate tissue patterns, early bottlenecks and lineage commitments lead to substantial variation in embryonic patterns both within and between individuals.

## INTRODUCTION

All cells in the adult human derive from a single fertilised egg following a well-orchestrated procession of cell division, cell movement and cell differentiation during embryonic and foetal development which continues throughout life. Tracing the lineages of cells can elucidate these fundamental developmental processes and has been widely employed in model organisms. Early lineage tracing experiments relied on forms of light microscopy^1^, an approach still used for studying early embryogenesis^2,3^. However, this becomes impractical over long periods of time and for complex organisms. Genetic tagging approaches^4,5^ allow detection of introduced markers at later points in life, but rely on a discrete period of genome scarring during development and their invasiveness obviously precludes their application to human development.

From fertilisation onwards, the cells of the human body continuously experience damage to their genome, either from intrinsic causes or from exposure to mutagens^6^. Although almost all DNA damage is repaired and the genome is replicated with extremely high fidelity, cells steadily acquire somatic mutations throughout life. Any cell, therefore, possesses a near-unique set of mutations. Some mutations in this set, however, are shared with other cells and such mutations indicate that a developmental history is shared by this group of cells, all of which are likely to be progeny of the cell that originally acquired the mutation. Once acquired, mutations are rarely lost. Thus, using acquired mutations, past developmental relationships from embryonic, foetal, childhood and adult phases of life can be identified from adult cells^7–11^ (Park et al, 2020).

The most direct way to reconstruct phylogenies of human development requires readouts of somatic mutations from single cells. This is possible through single-cell genomics^12,13^, but this can suffer from high rates of genomic dropout and large numbers of artefacts introduced during whole-genome amplification. Alternatively, single stem cells can be expanded *in vitro* into colonies or organoids and be subjected to standard whole-genome sequencing (WGS)^7,10^. However, information concerning the spatial relationship between different cells or clones is usually lost using this approach. Recently, low-input WGS following laser-capture microdissection (LCM) has allowed reliable calling of somatic mutations in distinct single-cell derived physiological units, such as colonic crypts^14^ and endometrial glands^15^, while retaining spatial information on a microscopic level. Here, we reconstruct large-scale phylogenies of cells from many organs from three individuals, reflecting tissue development and maintenance from zygote until death. The use of LCM allows us to directly combine genetic and spatial information to elucidate the embryonic origins and mosaic patterning of tissues (**Fig. 1a**).

**Figure 1.**
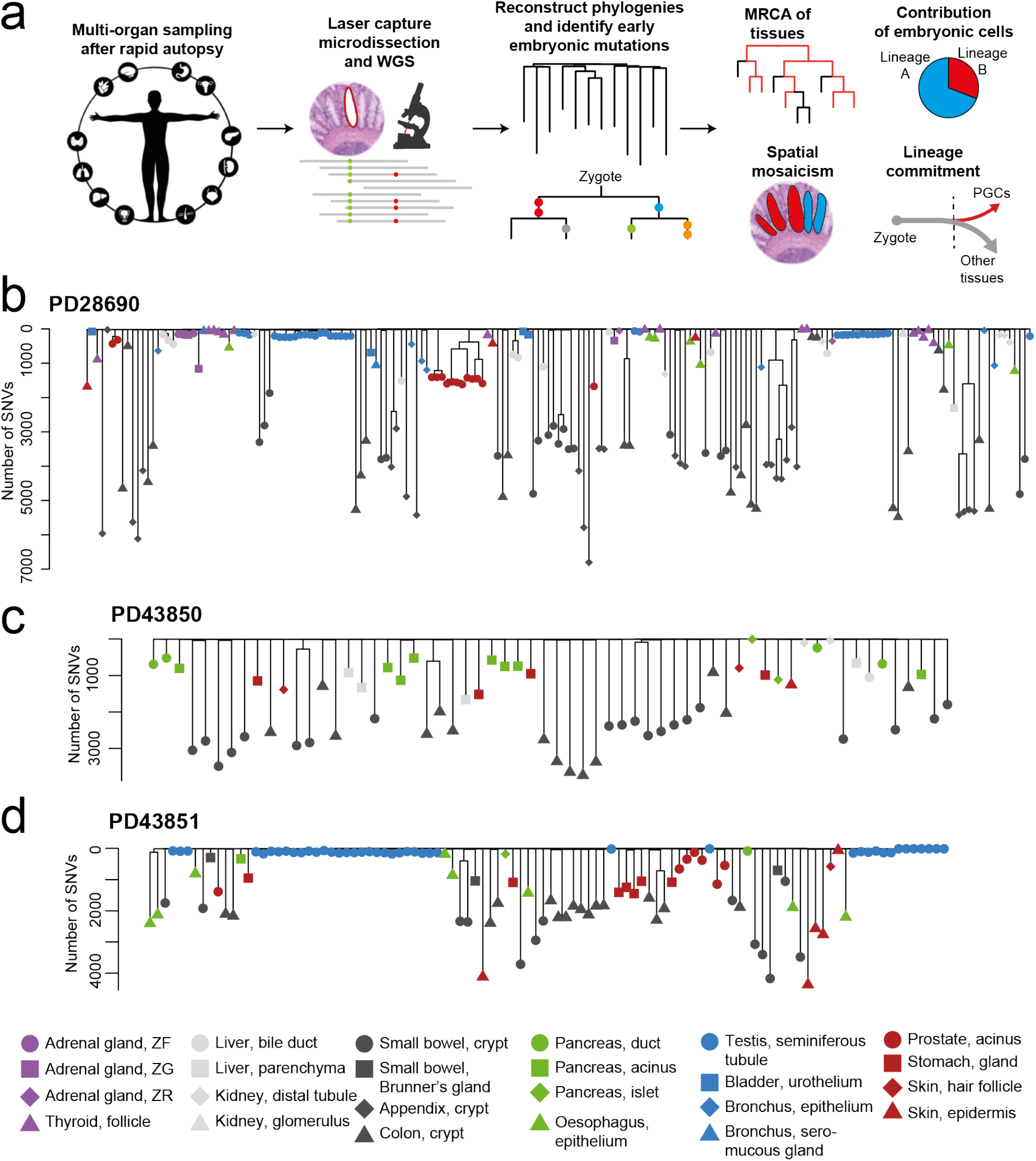
Phylogenies of clonal populations. Overview of experimental design and analysis. PGCs, primordial germ cells. Phylogenetic trees of clonal LCM cuts of PD28690 (**b**), PD43850 (**c**) and PD43851 (**d**). The branch length conveys the number of SNVs acquired in that lineage. The shape and colour of the dots reflect the tissue type of the sample as per the legend. ZF, zona fasciculata; ZG, zona glomerulosa; ZR, zona reticularis.

## RESULTS

### Phylogenies of normal tissues

We studied three individuals all of whom had been subject to autopsy shortly after death, PD28690 (male, 79 years old), PD43850 (female, 51), and PD43851 (male, 47). A total of 335 LCM dissections (each consisting of approximately 200-1000 cells) from 25 different tissues from PD28690, 66 from 12 tissues from PD43850 and 110 from 12 tissues from PD43851 were subjected to low-input WGS (**Fig. 1a**, **Extended Data Table 1**). We called single nucleotide variants (SNVs) against the human reference genome. Germline and artefactual variants were filtered out using an exact binomial test and a test for beta-binomial over-dispersion^8,16–18^ (**Methods**).

To directly reconstruct a phylogeny, only somatic mutations from single cell-derived clones were included. The clonal organisation of most of the tissue units sequenced here was not known prior to this experiment. However, the distribution of the variant allele frequencies (VAFs) of acquired mutations in each microbiopsy informs about the clonality of the cell population. For instance, the population of cells in a colonic crypt derive from a single stem cell at the crypt base which translates into a VAF distribution of heterozygous mutations centred around 0.5.^14^ Using a binomial mixture model solved by expectation-maximisation we estimated the number of clones and their contribution in each microdissection. This sorted the data into (mono)clonal, oligoclonal and polyclonal samples (**Extended Data Fig. 1**). To include a microdissection in the phylogeny reconstruction, a clone needed to account for the majority of cells.

Using sufficiently clonal samples we constructed phylogenies based on maximum parsimony and mapped SNVs onto the branches (**Fig. 1; Methods**). In total, 187 clonal samples were used for the phylogeny reconstruction of PD28690, 62 for PD43850, and 106 for PD43851. When branch lengths represent the number of SNVs, all trees exhibit a “comb-like” structure, revealing that most genetic sharing occurs in a short amount of time at the top of the tree, after which most SNVs are acquired privately by the lineage of cells leading to each dissected sample. The terminal branch length reflects the mutation burden of the last single cell ancestor generating an individual microdissected cell population, which may have existed many years prior to sampling. Intestinal crypts exhibit a high burden, while dissections of seminiferous tubules exhibit a much lower mutation burden (Moore et al, 2020).

### Embryonic dynamics and asymmetric contributions

The topologies of the phylogenetic trees reveal the pattern of branching events during development, mostly during the first days of embryogenesis (**Fig. 2a-c**). There are important qualifications in the interpretation of these trees. First, while every internal node depicts an ancestral embryonic cell, not all early cells are represented as nodes in the phylogeny. Due to cell death or commitment to extraembryonic lineages, embryonic cells may be cryptically present in branches or absent from the phylogeny altogether. Second, not all cell divisions can be resolved into bifurcations. If no somatic mutations occur at a division it becomes invisible and the resulting lineages are resolved into multifurcations (polytomies).

**Figure 2.**
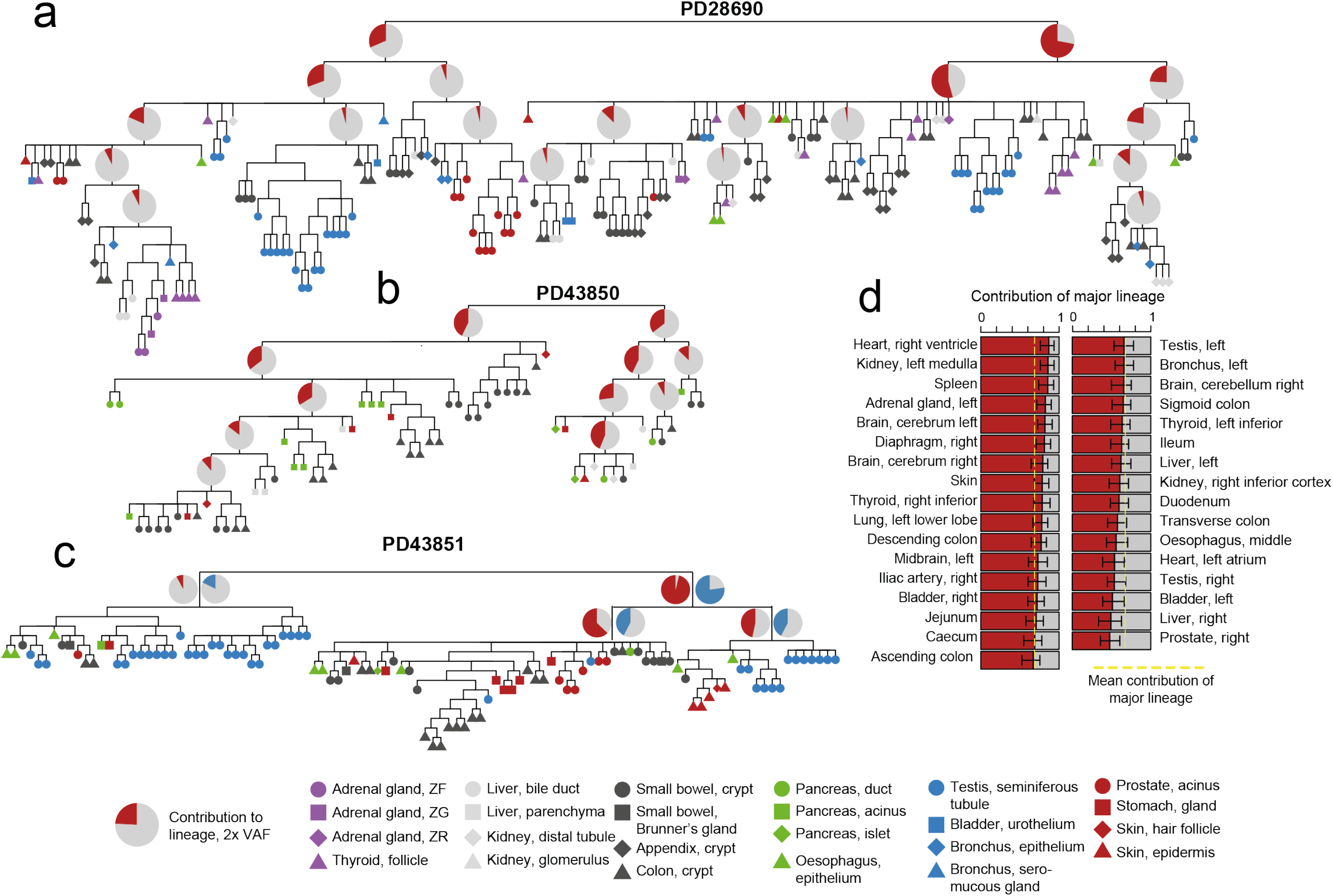
Developmental phylogenies and embryonic asymmetries Phylogenetic trees for PD28690. (**a**), PD43850 (**b**) and PD43851 (**c**). The shape and colour of the labels indicate tissue type as per the legend. Pie charts show the mean contribution (twice the VAF) of that lineage to the 33 bulk biopsies of PD28690, to a bulk brain sample of PD43850, or a bulk brain (red) or colon (blue) sample of PD43851. ZF, zona fasciculata; ZG, zona glomerulosa; ZR, zona reticularis. (**d**) The contribution of the major lineage to each bulk biopsy in PD28690, with 95% confidence intervals. The mean contribution of the major lineage is shown with the yellow dotted line.

In all cases, the tree starts in a bifurcating fashion for roughly two generations before preferentially generating large multifurcations. This is consistent with a higher mutation rate in the first few divisions of life, followed by a sharp decrease, which may be due to zygotic genome activation and consequent acquisition of full DNA repair capacity, occurring after a few cell divisions.^19^ It is also possible, however, that the pattern is due to more cell death or contribution to extraembryonic tissues during the earliest cell divisions and thus inclusion of more cryptic cell divisions in very early branches. We estimate a mutation rate per generation during the first two generations of 2.4 (95% confidence interval: 1.7-3.2), which drops to 0.7 (95% confidence interval: 0.51-0.77) during subsequent generations (**Methods**).

In PD28690, the most recent common ancestor for all individual tissue types is the root of the tree, presumably the zygote (**Extended Data Fig. 2**). The commonality of the most recent common ancestor indicates that no tissue studied here has a post-zygotic monophyletic origin beyond the cell giving rise to the entirety of the embryo. This polyphyletic origin for all tissues is in line with lineage commitment occurring during later stages of embryogenesis with a much larger cell population. The same overall pattern is present in the phylogenetic trees of PD43850 and PD43851.

We previously observed an asymmetric contribution of the two daughter nodes of the presumed zygote, such that one contributes twice as much to the adult body than the other^7,9,10^. It has been postulated that this is a consequence of differential cell allocation to the trophectoderm and inner cell mass during blastulation. The contribution of early embryonic progenitors can be measured through the VAF of the mutations on branches defining them. Bulk samples, i.e. large pieces of tissues, represent polyclonal aggregates of cells and should give an adequate population-level genomic readout of embryonic ancestry.

For PD28690, 33 bulk WGS samples from 18 different organs were used to assess the body-wide contribution of embryonic progenitors. The first observed bifurcation exhibits a mean VAF of 0.36 in the major lineage and 0.16 in the minor lineage, corresponding to an asymmetry of 69:31 (**Fig. 2a**). Since the sum of VAFs of these two lineages approximate 0.5, no appreciable cell lineages are missing from the first bifurcation (**Extended Data Fig. 3**). This asymmetry is reflected in most individual bulk samples as well, meaning that all are derived from a large population of ancestral cells (**Fig. 2d**). There is an establishment of further asymmetries in subsequent generations. The major lineage splits into two unequally contributing lineages with mean VAFs of 0.27 and 0.12, while the minor lineage splits in 0.15 and 0.02.

For PD43850, we estimate early lineage contribution using a single polyclonal bulk sample derived from brain, an organ unrepresented in the phylogeny. PD43850 exhibits a modest asymmetry compared to PD28690 (56:44, **Fig. 2b, Extended Data Fig. 3**), although this could be due to the paucity of available bulk samples.

For PD43851, a bulk sample from both brain and colon were available. In the primarily ectoderm-derived brain sample, the stark asymmetry (92:8) indicates it was almost entirely derived from one of the two first lineages (**Fig. 2c, Extended Data Fig. 3**). However, in colon, the asymmetry amounts to 81:19, significantly different from the observed asymmetry in brain (p=0.003, likelihood ratio test, **Methods**). Besides brain, the only other ectodermal tissue included in the phylogeny of this patient are microdissections of epidermis, which are exclusively derived from the major lineage. Interestingly, the minor lineage is ancestral to the large majority of seminiferous tubule microdissections, most from a distinct daughter lineage, and their distribution (35:65) is significantly different from the asymmetry in brain and colon (p<10^−8^ for all comparisons, binomial test). However, endoderm-derived microdissections are distributed on the phylogeny (81:19) in line with the asymmetry in colon (p=0.86), but not brain (p=0.005). Subsequently, the first split in the major lineage of the brain exhibits a similar asymmetry (64:36) as observed in PD28690, while this amounts to 50:50 in the colon.

Taken together, the observed asymmetric contributions of the first few cell generations can exhibit a similar pattern across all tissues in some individuals, such as PD28690, while revealing stark contrasts between different lineages in others, such as PD43851. In essence, the asymmetry of the first embryonic bifurcation is distinctly variable in humans and its apparent tissue-specificity suggests the existence of strong bottlenecks beyond blastulation.

### Clonal patterns and microscopic mosaicism

Next, we combined the genetic information from WGS with the spatial information from histological sections to reconstruct the mosaic patterns of cells laid out early in development. The epithelium of the intestine is organised as a two-dimensional array of crypts oriented perpendicular to the luminal surface. Results from jejunum and ileum (**Fig. 3a**) show that adjacent and nearby crypts usually arise from the same embryonic progenitor. The most recent common ancestor of such related crypts can be timed by the mutation burden of their ancestral lineage and indicated that they almost always existed during embryogenesis, rather than later in life. Thus during embryonic development, single cells seed patches of contiguous gut epithelium and the topology of these patches is still represented in the adult intestine. Sometimes, however, nearby crypts were from a different embryonic lineage, presumably representing an adjacent embryonic patch.

**Figure 3.**
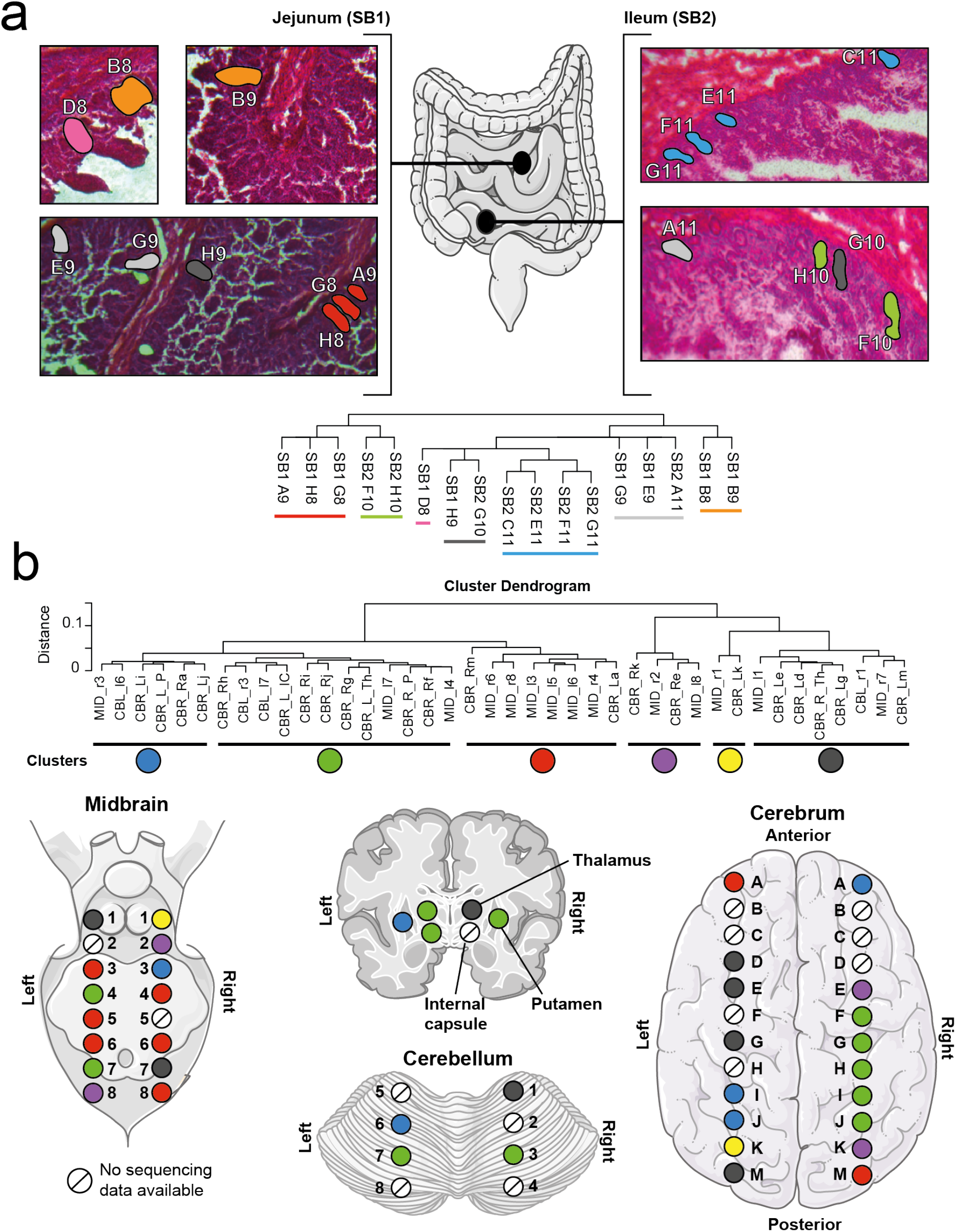
Embryonic mosaicism in tissues and organs. Histology sections of ileum and jejunum of PD28690 coloured by location on the phylogeny. (**b)** Cluster dendrogram of soft cosine similarity of VAFs of early embryonic mutations in bulk brain samples of PD28690. The data is split into six clusters, which are spatially displayed in the various brain regions. CBL, cerebellum; CBR, cerebrum; IC, internal capsule; L, left; MID, midbrain; P, putamen; R, right; Th thalamus. Parts of the figure are composed of pictures from Servier Medical Art. Servier Medical Art by Servier is licensed under a Creative Commons Attribution 3.0 Unported License.

In general, later clonal expansions are mostly confined to a few notable events in seminiferous tubules, colonic or appendiceal crypts (**Extended Data Fig. 4**). These expansions are often accompanied by driver mutations in canonical cancer genes, such as *APC*, *BRAF* or *GNAS*. The notable exception is an extensive expansion in the prostate of PD28690, consistent with benign prostatic hyperplasia in the absence of a discernible driver mutation. In all other cases, the genetic relationship between microdissections of the same biopsy is limited to that acquired during early development.

Although highly ordered tissues with distinct microscopic structures, such as intestinal epithelium, lend themselves to this type of analysis, echoes of embryonic patterning can be seen in other adult tissues. For example, microdissections of epidermis often yield polyclonal samples, which can be decomposed into their embryonic lineages using the VAFs of mutations in the phylogeny. The VAF patterns in epidermal microdissections of PD28690 reveal that some are aggregates of multiple embryonic clones and represent a mixing of different seeding events (**Extended Data Fig. 5**). Others manifest as a single clonal lineage in the early embryo, indicating the existence of a single developmental founder cell, the progeny of which dominates this sample. The low mutation burden and exclusion from the phylogeny construction indicates this clonal phase is very short, and hence cannot represent a later clonal expansion.

### Organ-wide macroscopic mosaicism in the brain

The profile of VAFs of embryonic mutations in an individual bulk sample is a summation of the developmental history, from the time each mutation arose to the time of sampling, of every cell in the sample. It will reflect the aggregation of all the constituent founder cell types and developmental bottlenecks, recent and in the more distant past of the sample population. This will be the case even if the mutations are acquired prior to lineage commitment.

To assess the broad spatial patterning laid out in organ development, 86 different bulk samples of PD28690 (including 40 samples derived from various regions of the brain) were subjected to high-depth targeted re-sequencing of embryonic variants. Samples were then clustered together based on the soft cosine similarity of the VAF profiles of embryonic mutations (**Methods**). Brain samples fell into roughly six different clusters (**Fig. 3b**). Strikingly, adjacent biopsies in the cerebrum and midbrain often belong to the same cluster. It is plausible that this is a consequence of a shared developmental path and the seeding of those areas by similar populations of cells. Moreover, the different regions of the brain show a heterogeneity in the prevalence of different clusters, with one particular cluster being prominently present in the midbrain. The correlation between the spatial and genetic distance was significant in the cerebrum but not the midbrain (p=0.039 and p=0.27, Mantel test). This means that, despite these bulk samples being taken millimetres apart and being composed of many different cell types, there is greater sharing of a specific VAF pattern of pre-gastrulation mutations between neighbouring regions of the cerebrum compared to distant regions. Taken together, this data indicates that developmental bottlenecks and lineage commitments generate a spatial effect in the mosaicism of the brain, especially in the cerebrum.

### Separation of primordial germ cells from somatic lineages

Prior to or around gastrulation, the lineage of primordial germ cells (PGCs) is thought to be the first to completely segregate from other lineages^20^. Embryonic mutations found in descendants of PGCs that occurred before this segregation may therefore be seen in tissues derived from the three major germ layers, whereas those arising after this segregation should not be. The descendants of PGCs in men include spermatogonia, the adult stem cells in the seminiferous tubules of the testis. Spermatogonia and their descendants account for most cells in microdissections of individual seminiferous tubule cross sections, which are often monoclonal. In PD28690 and PD43851, seminiferous tubule microdissections shared on average 7.0 and 8.7 SNVs respectively with any other tissue microdissection. The observed segregation of PGCs in the phylogeny of PD28690 was confirmed by spermatogonia-specific variants (**Extended Data Fig. 6**). Further sequence data from 162 microdissections of seminiferous tubules from 11 individuals from whom colon or blood samples were also available (**Extended Data Table 1**) showed that, on average, seminiferous tubules shared 4.5 mutations with the bulk sample (**Fig. 4a**). This is consistent with PGC specification occurring prior to gastrulation. Furthermore, some seminiferous tubules shared no SNVs whatsoever with their matched bulk sample. In PD40745, the root of the phylogeny splits into three lineages of seminiferous tubules with only two making a detectable contribution to bulk **colon**. This suggests that, due to later embryonic bottlenecks, a subset of PGCs genetically segregated from the cells generating the three major germ layers after the first cell divisions of life.

**Figure 4.**
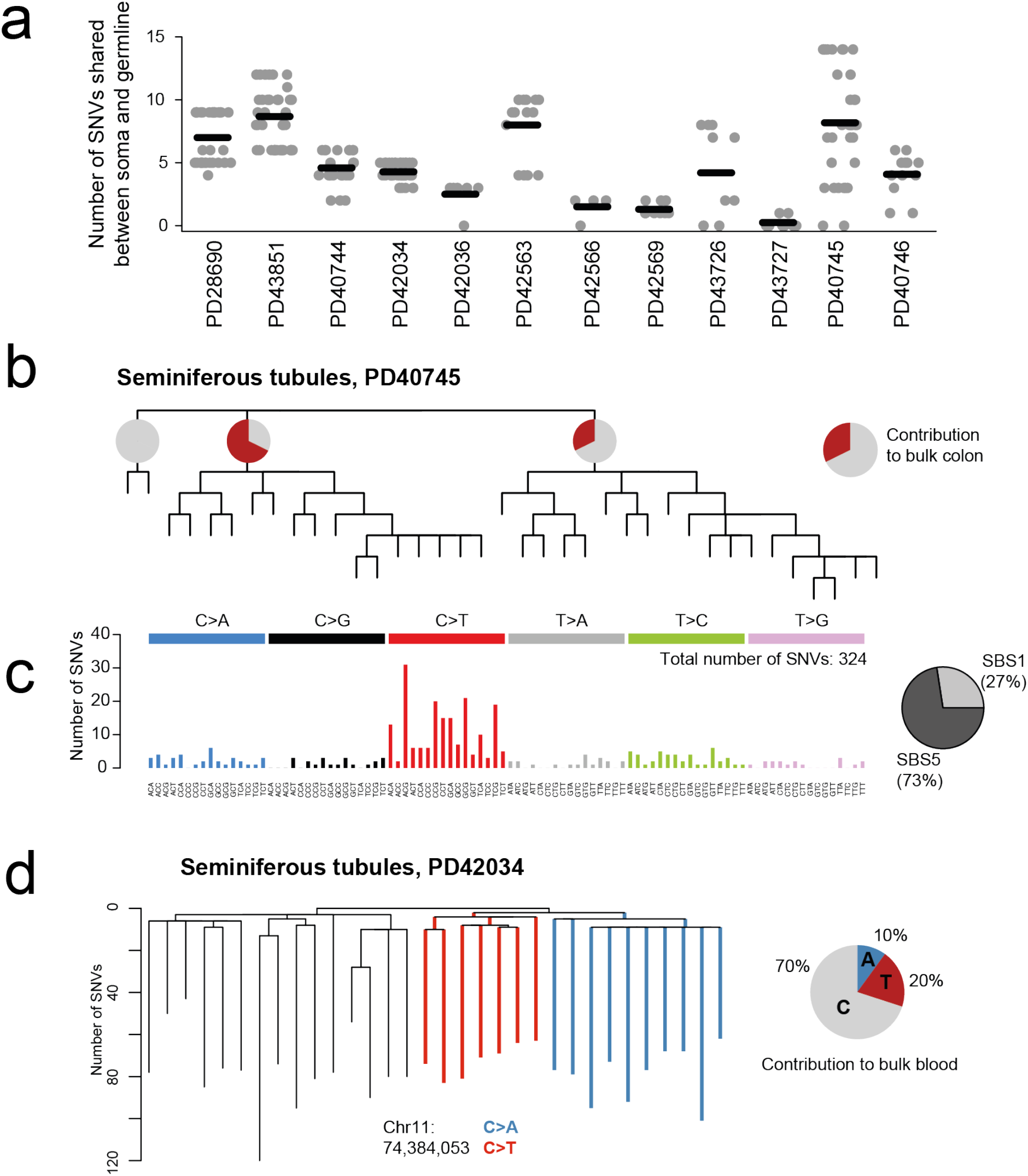
Patterns of mutations in early embryogenesis. **(a**) Number of SNVs shared between any seminiferous tubule microdissection and another microdissection in PD28690 and PD43851. In other patients, number of SNVs shared between seminiferous tubule microdissections and matched bulk sample. (**b**) Phylogeny of seminiferous tubules of PD40735, with early lineage contribution to bulk colon in pie charts, showing a lineage of seminiferous tubules undetectable in bulk colon. (**c**) Mutational spectrum of early embryonic SNVs. **(d)**SNV occupying the same site in the genome in sister clades. Pie chart depicts the contribution (twice the VAF of SNVs) to matched bulk colon.

### Patterns of mutations and recurrent LOY

Embryonic SNVs, as characterised by their trinucleotide context, exhibit a mutational spectrum consistent with mutational processes ubiquitous in normal tissues (**Fig 4c**., **Methods**, Moore et al., 2020). In PD43851, two different SNVs occupy the same site in large neighbouring clades of seminiferous tubules, a C>A and a C>T mutation, both of which are detectable in bulk blood (**Fig. 4d**). It is possible that this is a consequence of an unrepaired DNA lesion passed on after the first observed cell division in the phylogeny, which is subsequently repaired differentially in the two daughter clades. This would be an *in vivo* observation in normal tissues of DNA lesion segregation.^21^

Other types of somatic mutations, such as insertions and deletions (indels), copy number variants (CNVs) and structural variants (SVs) occur at a much lower rate than SNVs and are therefore of limited use for phylogeny reconstruction. The only CNV affecting multiple tissues was a loss of the Y chromosome (LOY). Its distribution on the phylogeny indicates a recurrent late event rather than a single early event (**Extended Data Fig. 6**). In PD28690, this affected around 6% of all LCM cuts. No LOY was detected in samples from the other male patient, PD43851, nor any of the seminiferous tubules from 11 patients. LOY affected multiple samples of bladder urothelium, bile duct, adrenal gland and renal distal tubules in PD28690, suggesting a tissue-specific propensity for LOY. This tissue-specificity mirrors the one reported for different tumour types^22^. LOY has been associated with a wide variety of age-related disorders and may act as a proxy for genomic instability^23–25^.

In contrast to reports on widespread aneuploidy in pre-implantation human embryos^26–28^, we observed no early SVs or CNVs in any patients studied. This may suggest that aneuploid cells are purified from the lineages constituting the embryo proper, either through cell death or allocation to extraembryonic lineages.

## DISCUSSION

This research exemplifies the use of somatic mutations as markers for lineage tracing in human development. By constructing phylogenetic trees from samples of many normal tissues, it is possible to follow the life history of a cell from its first beginning as a zygote, through embryogenesis and early development, all the way to its final destination as a differentiated adult cell.

The widespread variability in observed asymmetry in embryonic contribution suggests that the precise lineage commitment of cells is not fixed. Unobserved effects, such as the spatial positioning of individual cells in the early embryo, might be at the heart of creating seemingly stochastic patterns of asymmetries while resulting in the same developmental outcome. While the early segregation of trophectoderm and inner cell mass might explain the global asymmetry in PD28690, later bottlenecks can tune the levels of asymmetry manifesting in individual tissues and lineages, as seen in PD43851. To be detectable at a resolution of bulk samples, these bottlenecks must be strong and reflect that many lineages derive from small subsets of cells during development. After blastulation, the most notable developmental bottlenecks or lineage commitments will be the differentiation between epiblast and hypoblast, PGC commitment^20^, gastrulation, extraembryonic intercalatation^29,30^ and organogenesis. These later bottlenecks may have resulted in the strong spatial effect on genetic mosaicism in the cerebrum of PD28690.

Somatic mutations enable the tracing of individual lineages and the timing of their segregation, leading to the observation of an early, pre-gastrulation segregation of the PGC lineage. As an extension to this, studying patterns of later lineage commitments, such as those occurring in gastrulation, must rely on phylogeny reconstruction of a larger number of samples, to exhaustively sample the cell population at the time of commitment. The resolution can be further enhanced by high-depth targeted resequencing of populations of pure cell types.

Understanding the patterns of normal human development can contextualise disorders of development, especially when caused by post-zygotic genomic aberrations, such as early drivers of childhood cancers or mosaic overgrowth syndromes.^8^

Taken together, this research demonstrates the variability of human embryogenesis in the role and prominence of individual embryonic progenitors and the shaping of different lineages and tissues through developmental bottlenecks. Further studies employing somatic mutations as natural lineages markers can help answer fundamental questions of human ontogeny.

## METHODS

### Ethics statement

Metadata on all samples can be found in **Extended Data Table 1**.

- **Donor PD28690**: Multiple samples from 22 macroscopically normal tissues and organs (Supplementary Table 1) were collected from a 78-year-old male during a rapid autopsy (rapid autopsy defined as an autopsy with a post-mortem time interval (PMI) of less than six hours). This donor was a non-smoker who died of a metastatic oesophageal adenocarcinoma for which he had received a short course of palliative chemotherapy (5-6 weeks of oxaliplatin 7 weeks prior to death). He had no other comorbidities. The samples were collected in line with the protocols approved by the NRES Committee East of England (NHS National Research Ethics Service reference 13/EE/0043). Every sampled tissue was photographed and biopsy sites documented. Once collected, all tissue biopsies were snap frozen in liquid nitrogen and subsequently stored at −80°C.
- **Donors PD43850 and PD43851:** Multiple biopsies from 16 different tissues were collected from a 54 year-old female (PD43850) and a 47 year-old male (PD43851); both individuals died of non-cancer causes (traumatic injuries and acute coronary syndrome respectively). Similarly, all samples were obtained within less than six hours of death (one hour and three hours respectively), were snap frozen in liquid nitrogen and subsequently stored at −80°C. The use of these tissues was approved by the London, Surrey Research Ethics Committee (REC reference 17/LO/1801, 26/10/2017).
- **Additional testis and colon samples:** AmsBio (commercial supplier) – Samples for PD40744, PD40745, PD40746, PD42563, PD42566 and PD42569 were obtained at autopsy from individuals who died on non-cancer related causes. The use of these tissues was approved by the London, Surrey Research Ethics Committee (REC reference 17/LO/1801, 26/10/2017). Samples for PD42036, PD42034, PD43727, PD43726 and PD46269 were obtained from individuals who had non-testicular problems such as abdominal chronic pain and tissue distortion. The use of these samples was approved by North East Newcastle & North Tyneside1 (REC reference 12/NE/0395, 22/09/2009). As above, all collected samples were snap frozen in liquid nitrogen and subsequently stored at −80°C.

### Bulk DNA sequencing

DNA was extracted from bulk biopsies, short insert (500bp) genomic libraries were constructed, flow cells prepared and 150 base pair paired-end sequencing clusters generated on the Illumina HiSeq X Ten platform according to Illumina no-PCR library protocols. An overview of samples, including the average sequence coverage, is shown in Extended Data Table 1.

### Laser capture microdissection and low-input DNA sequencing

Tissues were prepared for microdissection and libraries were constructed using enzymatic fragmentation as described previously^14,15^ and subsequently submitted for whole-genome sequencing on the Illumina HiSeq X Ten platform. An overview of samples, including the average sequence coverage, is shown in **Extended Data Table 1**.

### DNA sequence alignment

All DNA sequences were aligned to the GRCh37d5 reference genome by the Burrows-Wheeler algorithm (BWA-MEM).^31^

### Somatic variant calling

Single nucleotide variants (SNVs) were called using the CaVEMan (Cancer Variants through Expectation Maximisation) algorithm, a naïve Bayesian classifier.^32^ Traditionally, it is used to call somatic variants in one sample, usually tumour, by using another sample from the same patient as a matched normal. However, to preserve the early mutations that will be present in normal bulk samples, we used an unmatched normal sample. This alignment file was created *in silico* by generating reads from the human reference genome (GRCh37). In effect, this causes all nucleotide deviations with respect to the reference genome to be reported, including a large amount of inherited single nucleotide polymorphisms (SNPs). However, CaVEMan automatically employs its flags and filters when calling variants, even in an unmatched setting. One of such filters is the exclusion of a putative SNV that is present in a large panel of normal samples. This excludes most of the germline SNPs from subsequent analysis and brings their total number down from approximately 4-5 million to 30,000-40,000 in most unmatched runs. The latter represents rare inherited SNPs. In addition to the default CaVEMan filters, putative SNVs were forced to have a mean mapping score (ASMD) of at least 140 and fewer than half supporting reads being clipped (CLPM=0).

Where whole-genome sequencing data was obtained from LCM samples through the low-input pipeline, an additional set of filters was used to remove specific artefacts, as previously described.^14,15^ These artefacts include spurious variant calls due to the formation of cruciform DNA and the double counting of variants due to a negative insert size.

Short insertions and deletions (indels) were called using the Pindel algorithm,^33^ with calling and filtering strategies the same as for SNVs. Copy number variants (CNVs) were called using the ASCAT (Allele-Specific Copy number Analysis of Tumours) algorithm^34^ and the Battenberg algorithm.^35^ Since both ASCAT and Battenberg rely on the sharedness of SNPs, they cannot be run against the *in silico*-generated unmatched normal sample. To detect potential early embryonic CNVs, both ASCAT and Battenberg were run on all LCM samples against bulk samples and alternatively, against an LCM sample known to be derived from the other branch of the first split in the phylogeny, and hence genetically unrelated on a post-zygotic level. No early embryonic CNVs were present in any of the phylogenies.

To detect structural variants (SVs) the BRASS (BReakpoint AnalySiS) algorithm^35^ was used, which relies on discovery through discordantly mapped reads and confirmation by local reassembly of the breakpoints of the SV. The strategy for matching samples was identical to the strategy described for ASCAT and Battenberg. Again, no early SVs were identified in any of the phylogenies.

### Filtering shared artefacts

Given that inherited variants are expected to be present at least at a heterozygous level in all cells, a one-sided exact binomial test can be used on the aggregated counts of reads supporting the variant and the total depth at that site. Basically, this tests whether the observed variant counts are likely to have come from a germline distribution (given the total depth), or whether it is more probable to come from a distribution with a lower true VAF. For sex chromosomes in male patients, the binomial probability (true VAF) for comparison was set to 0.95 rather than 0.5. This results in a probability per variant whether it is inherited. These values are corrected for multiple testing using a Benjamini-Hochberg correction.^36^ Any variant with a corrected p-value less than 10^-5^ is categorised as a putative somatic variant.

### Filtering shared artefacts

To filter out recurrent artefacts, a beta-binomial distribution is fitted to the number of variant supporting reads and total number of reads across samples from the same patient. For every somatic SNV, maximum likelihood over-dispersion parameter (σ) is determined in a grid-based way (ranging the value of σ from 10^-6^ to 10^-0.05^). A low over-dispersion captures artefactual variants, as they appear seemingly randomly across samples and can be modelled as drawn from a binomial distribution. In contrast, true somatic variants will be present at a high VAF in some, but not all genomes, and are thus best represented by a beta-binomial distribution with a high over-dispersion. To distinguish artefacts from true variants, a threshold was set at σ=0.1, below which variants were considered to be artefacts. The code for this filtering approach is an adaptation of the Shearwater variant caller.^37^

While the number of variants that are filtered out in this step is usually rather modest (in most cases this will be fewer than a 1,000 mutations), it greatly enhances the ability to reconstruct genuine phylogenies of the samples. This over-dispersion filter will remove any inherited mutations that have falsely passed the filters so far (as they will be completely under-dispersed) and also removes recurrent artefacts that might wrongly be interpreted as mutations occurring early in development.

### Truncated binomial mixture modelling

It is often the case with variant calling algorithms that there is a threshold for the number of reads supporting a variant before it is taken into consideration. This is mostly because of practical reasons: variants with low support will be abundant in the genome due to sequencing errors and noise in alignments leading to an increased run time of the algorithm whole increasing the rate of artefacts. For CaVEMan, the minimum support for a variant is hard-coded to be four reads.

This minimum number of supporting reads poses a problem when using a binomial mixture model on variants from a sample. In effect, part of the full binomial model will be censored, as observations from the lower tail of the distribution are disallowed. This will lead to an erroneous fitting of distributions, especially for clusters with a binomial probability approaching the edge of the censored distribution. The need for adjusted binomials also increases with lower depth of coverage, where the minimum threshold for support will more often be imposed on genuine variants. To fix this, truncated binomial distributions were used instead of full ones. In effect, this curtails the entire binomial probability distribution below a set threshold (here taken to be four reads), after which the remaining distribution is re-normalised.

### Estimating clonality

A mixture model can be applied when multiple different clones can inhabit the same sample, especially if the sample was a small section of tissue without much histological structure, such as skin or oesophageal epithelium. In this case, the model will try to separate the overall variants into the clones they could have arisen from, each with their own probability (VAF) and proportion (the amount of variants a clone contributes). Note that the latter proportion is the proportion of variants explainable by a clone, not the proportion of cells in a sample inhabiting that clone. The latter measure can be obtained by multiplying the VAF of a clone by two and is extremely useful to determine the clonal origin of any sample and whether two clones are nested or parallel. For example, clone A has an estimated VAF of 0.45 and clone B 0.15, so that the proportion of cells belonging to clone A is 90% and for clone B 30%. Simply because the sum of these proportions would exceed the total of the sample (120% is impossible), cells belonging to clone B must also belong to clone A. In other words, clone B must be a subclone of clone A. This logic is generally referred to as the pigeon hole principle.

Determining the largest clone in a given sample is straightforward with this framework. To reconstruct phylogenies we must be sure that our set of variants represent those coming from distinct single-cell derived clones and not mixtures that could be parallel i.e. sharing no genetic ancestry. As soon as any clone accounts for more than half of the cells in a sample, we can be sure that this signal is coming from a single ancestor, since two overlapping clones at 51% cannot co-exist. However, theoretically, as soon as a clone only accounts for (less than) half, it becomes possible for two clones to overlap, confuse the signal, and violate the assumptions of the phylogeny reconstruction. Therefore, only samples containing a clone with an estimated peak VAF higher than 0.25 are used for tree building.

In practice, many samples will have a degree of contamination from a polyclonal source, such as stroma. In such cases, the total clonal reconstruction will add up to less than 100%, because variants belonging to the polyclonal contaminant will have a true VAF of far below the detection limit in a typical whole-genome sequencing experiment. This contamination decreases the VAF of the largest clone and can therefore affect the estimated clonality of the sample. For example, with 20% stromal contamination, a clone comprising 60% of cells of interest will only account for 48% of the total sample, and hence, will not make the cut to be included in initial phylogeny reconstruction.

### Phylogeny reconstruction

Phylogenies of microdissected trophoblast clusters were generated from the filtered substitutions using a maximum parsimony algorithm MPBoot.^38^ MPBoot was run with default parameters (1,000 bootstrap iterations) on concatenated nucleotide sequences of variant sites. To ensure the proper anchoring of the root of the phylogeny, an artificial nucleotide sequence composed of the reference genome bases at variant sites was included to simulate the ancestral state at the zygote. Substitutions were mapped onto tree branches using a maximum likelihood approach. Any branch with a length of 0 after SNV fitting is collapsed into a polytomy.

In addition to the algorithms described above, the ape and ggtree packages were used for analysis and visualisation of phylogenetic trees in R.

### Mutation rate in early embryogenesis

The calculation of the mutation rate from the phylogenetic trees presented relies on the mutations per branch following a Poisson distribution, with the mutation rate itself as its only parameter (λ). The probability of observing a certain number of mutations in a branch (k) is then given by:

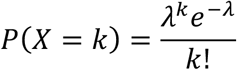

The rate parameter in a Poisson distribution is also the mean and the variance of the distribution. Given the absence of polytomies in the early branching events of the tree, the mutation rate can be calculated as the total number of mutations observed in the first two generations divided by the number of branches in those generations, the latter of which is equal to 6. Confidence intervals are then obtained using a Poisson test. This results in the estimates per patient, as follows:

|  | Number of SNVs | Rates | CI upper (95%) | CI lower (95%) |
| --- | --- | --- | --- | --- |
| PD28690 | 14 | 2.3 | 1.3 | 3.9 |
| PD43850 | 10 | 1.6 | 0.8 | 3.1 |
| PD43851 | 19 | 3.2 | 1.9 | 4.9 |
| Combined | 43 | 2.4 | 1.7 | 3.2 |

For the subsequent generations, where the occurrence of polytomies becomes so frequent, it is more straightforward to estimate the mutation rate by translating the observed number of branches involved in a multifurcation into the necessary frequency of branches with no mutations. The number of missed divisions is the number of branches involved in a polytomy minus two e.g. a trifurcation is the result of a single missed division; a four-way polytomy of two missed divisions and so on. Then, the following relation will hold:

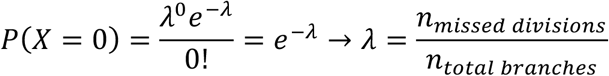

Following this, the calculations for the mutation rates of the third and fourth observed generation are:

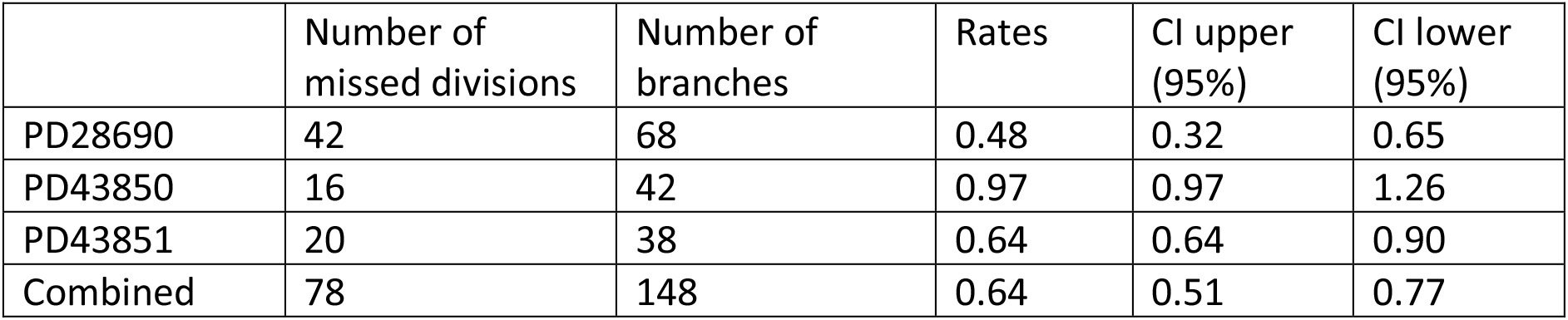

|  | Number of missed divisions | Number of branches | Rates | CI upper (95%) | CI lower (95%) |
| --- | --- | --- | --- | --- | --- |
| PD28690 | 42 | 68 | 0.48 | 0.32 | 0.65 |
| PD43850 | 16 | 42 | 0.97 | 0.97 | 1.26 |
| PD43851 | 20 | 38 | 0.64 | 0.64 | 0.90 |
| Combined | 78 | 148 | 0.64 | 0.51 | 0.77 |

### Likelihood ratio test

We used a likelihood ratio test to calculate whether two bulk samples have a significantly different asymmetry. The number of variant supporting reads (*N_V_*) and total reads (*NR*) for mutations on the major are modelled with a binomial distribution, with underlying probability *p*, while those on the minor branch have probability *1-p*. The likelihood of the model is then given as the binomial probability of seeing *N_V_* successes out of *N_R_* trials, given *p* (or *1-p*). The null model is that the underlying probability (asymmetry) is the same in the two bulk samples (*p1 = p2*), while the alternative model is that these are different (*p*_1_ ≠ *p*_2_). A grid-based approach was used to calculate the maximum likelihood estimates of the binomial probabilities.

### Targeted sequencing of embryonic variants

Following the identification of early embryonic variants in PD28690, a custom bait set for the Agilent SureSelectXT platform was designed using Agilent's online tool. Repetitive regions were masked using the criterion of least stringency, as defined by Agilent. Each embryonic variant was covered by one tile. In addition to early variants, testis-specific somatic mutations, as well as heterozygous SNPs were included in the bait set. The former was included to assess the split between the germline and the somatic tissues, while the latter was used to assess any biases in the performance of the pulldown. DNA libraries from 86 bulk samples were made and subsequently hybridised with the baits. The sequencing was performed on the Illumina HiSeq 2500 platform.

### Soft cosine similarity

To compare the similarity between two vectors of VAFs of variants positioned on a phylogenetic tree, a soft cosine similarity was calculated, which includes a specific similarity term to incorporate the dependence of observations. That is, variants on the same branch of the phylogeny will convey the same information, while variants of different branches of the tree will provide parallel information on lineage composition. As such, an interaction term *s_i,j_* is introduced into the following definition of the soft cosine similarity:

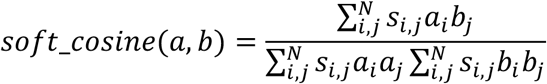

In this relation, *a* and *b* refer to two bulk samples and their vector of VAFs of embryonic mutations of length *N*, which are indexed by *i* and *j*. These VAFs are log-transformed. The similarity metric *s_i,j_* is defined as follows:

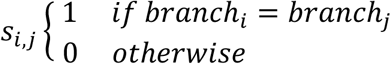

Note that if the similarity metric *si,j* is 1 for *i=j* and 0 otherwise, the formulation of the regular cosine similarity is retrieved.

To use the matrix of soft cosine similarities as a basis for clustering, it was converted to a soft cosine distance matrix by subtraction from 1 (i.e. if the soft cosine similarity between *a* and *b* is 0.7, the distance will be 0.3). Hierarchical clustering was then performed in R.

Significant correlations between the spatial and genetic distance matrices were assessed using the Mantel test, which is independent of the clustering.

### Embryonic SNVs and signature fitting

For the three patients subjected to a rapid autopsy, an embryonic SNV was defined as one present on a branch that was ancestral to two different tissue types. In the eleven men of whom primarily seminiferous tubules were samples, an embryonic SNV was defined as one shared between a seminiferous tubule and the matched bulk sample. Embryonic SNVs are listed in **Extended Data Table 2**.

Mutational signatures (COSMIC v3) were fitted using SigFit.^39^

### Recurrent SNVs and the infinite sites model

The assumption that somatic mutations generally represent unique events and do not happen twice is known as the infinite sites model and forms the cornerstone for maximum parsimony, the phylogenetic tree building method employed in this paper. The SNVs identified in this research and the constructed phylogenies can confirm or refute the validity of this assumption. One exception to the infinite sites model comes in the form of selected driver mutations, such as variants affecting hotspot regions in oncogenes. Indeed, in PD28690 we observe a recurrent mutation in *GNAS* (R844H) acquired independently in two appendiceal crypts.

In addition, a small number of SNVs in non-coding regions present as recurrent mutations in the same patient at the same site. This can be either the same nucleotide change i.e. a mutation violating the topology of the tree, or a different one. While the human genome is large, it is not infinite. This sequencing study catalogued over 100,000 SNVs in each individual studied. While this number is not yet sufficiently high to saturate the genome and invalidate all inferences on the timing of the acquisition of the SNV, it is enough to occasionally encounter independent SNVs at the same site or even the exact same SNV altogether. This probability can be quantified using a birthday paradox-inspired collision model. For ease of argument, we take the human genome to have three billion sites, all equally mutable. PD43850 and PD43851 each have slightly over 100,000 somatic mutations called, while PD28690 has approximately 300,000 in total. The probability of observing a site being mutated twice within the entire set of mutations is then 81% for PD43850 and PD43851, and approximately 100% for PD28690. If we require the exact same mutation to occur i.e. increase the number of possible mutations to nine billion, these probabilities become 43% and 99%, respectively.

It is important to note that this does not invalidate the overall maximum parsimony assumption nor the reconstructed phylogenies. Mutations occurring at the same site, but with different consequences, would be considered independent events. The occasional recurrent, independent mutations violating the phylogeny will be vastly outweighed by the numerous uniquely acquired mutations delineating human development. Hence, a low number of departures from the infinite sites assumption will not affect the overall tree topology. However, this might increasingly become a problem as the capacity for sequencing is increased and these lineage tracing efforts using somatic mutations are done at an even larger scale.

## ACKNOWLEDGEMENTS

We thank the staff of Wellcome Sanger Institute Sample Logistics, Genotyping, Pulldown, Sequencing and Informatics facilities for their contribution, especially Laura O’Neill, Calli Latimer and Kirsty Roberts for their support with sample management and laboratory work. We thank Sam Behjati, Young Seok Ju, Seongyeol Park, Jannat Ijaz, Pantelis Nicola and Grace Collord for helpful discussions or critical review of the manuscript.

## AUTHOR CONTRIBUTIONS

T.H.H.C., L.M., R.R., M.R.S. conceived the study design. T.H.H.C. wrote scripts and performed analyses with help or input from R.S., J.C., M.D.C.N. and I.M. L.M., P.S.R., A.C., and T.R.W.O. performed microdissections with support from Y.H. T.J.M., A.N. and R.F. aided in sample procurement. M.R.S. oversaw the study. T.H.H.C. and M.R.S. wrote the manuscript with input from all other authors.

**CORRESPONDING AUTHORS**

---

**Extended Data Figure 1.**
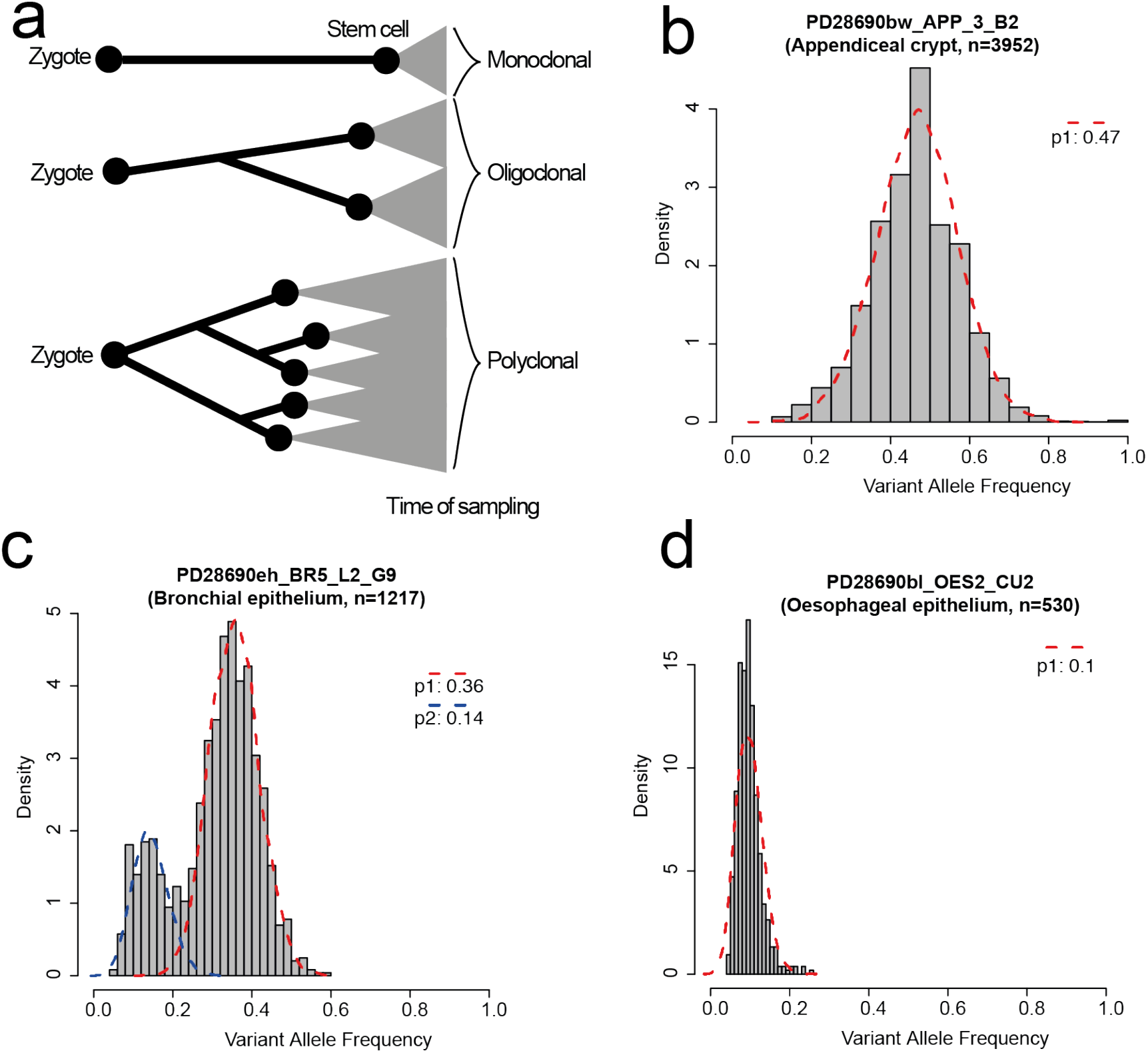
VAF distributions reflects clonality of LCM sample. **(a**) Schematic of three different progenitor or stem cell contributions to the eventual sample. Monoclonal samples consist of the progeny of one cell, while oligo- and polyclonal are derived from a few and many progenitors respectively. VAF histograms and binomial decompositions for a monoclonal (**b**), oligoclonal (**c**) and polyclonal (**d**) sample. Red and blue dashed lines indicate clonal decomposition through a binomial mixture model, with the estimated peak VAF of clones indicated in the legend. The number indicated in the title of each histogram is the SNV burden.

**Extended Data Figure 2.**
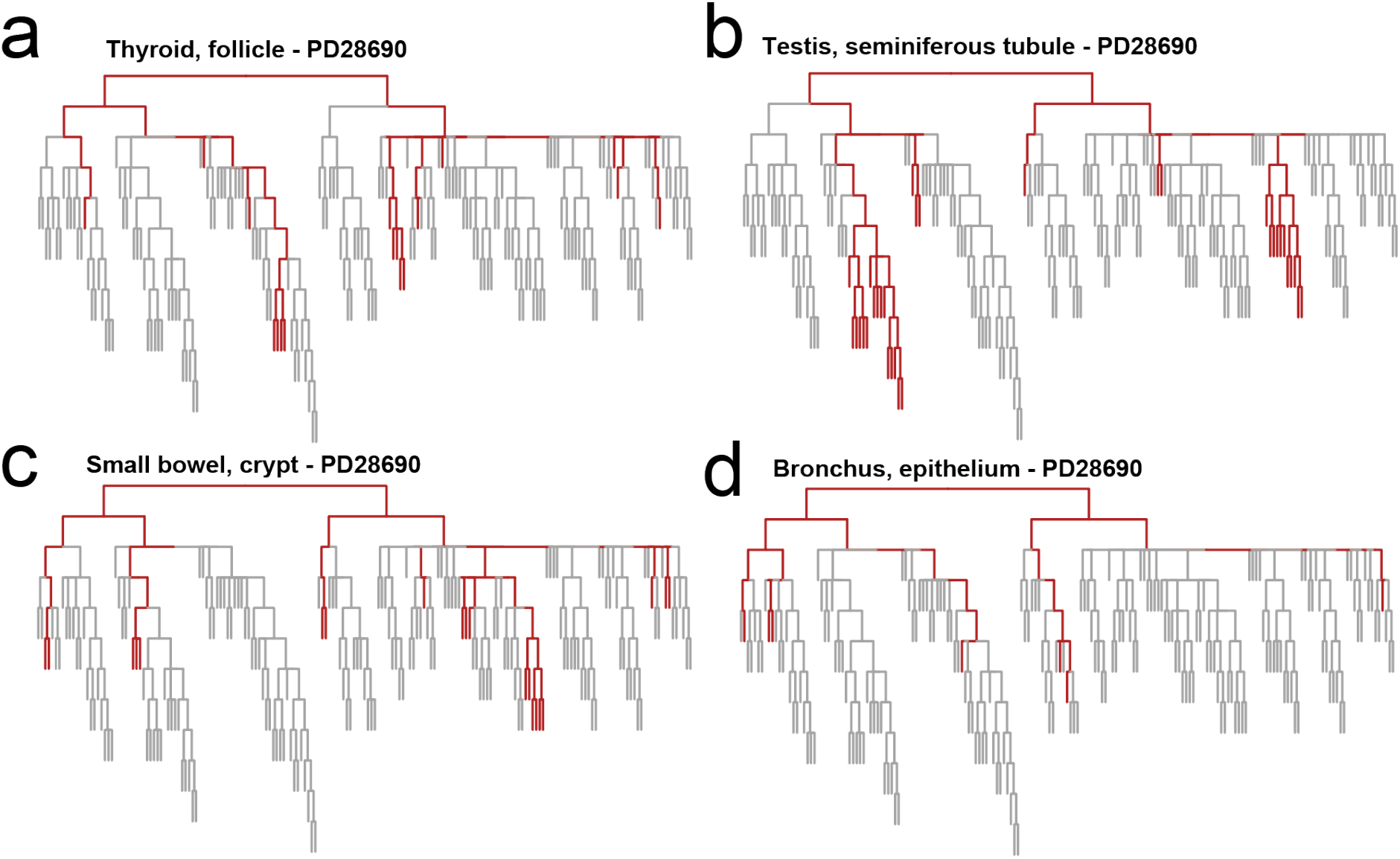
Most recent common ancestors of tissues. Phylogenetic trees with unit branch lengths for PD28690, showing the coalescence (red) of all samples from four tissues types: thyroid follicles (**a**), seminiferous tubules (**b**), small bowel crypts (**c**) and bronchial epithelium (**d**). The most recent common ancestor for all these tissues is the root of the tree.

**Extended Data Figure 3.**
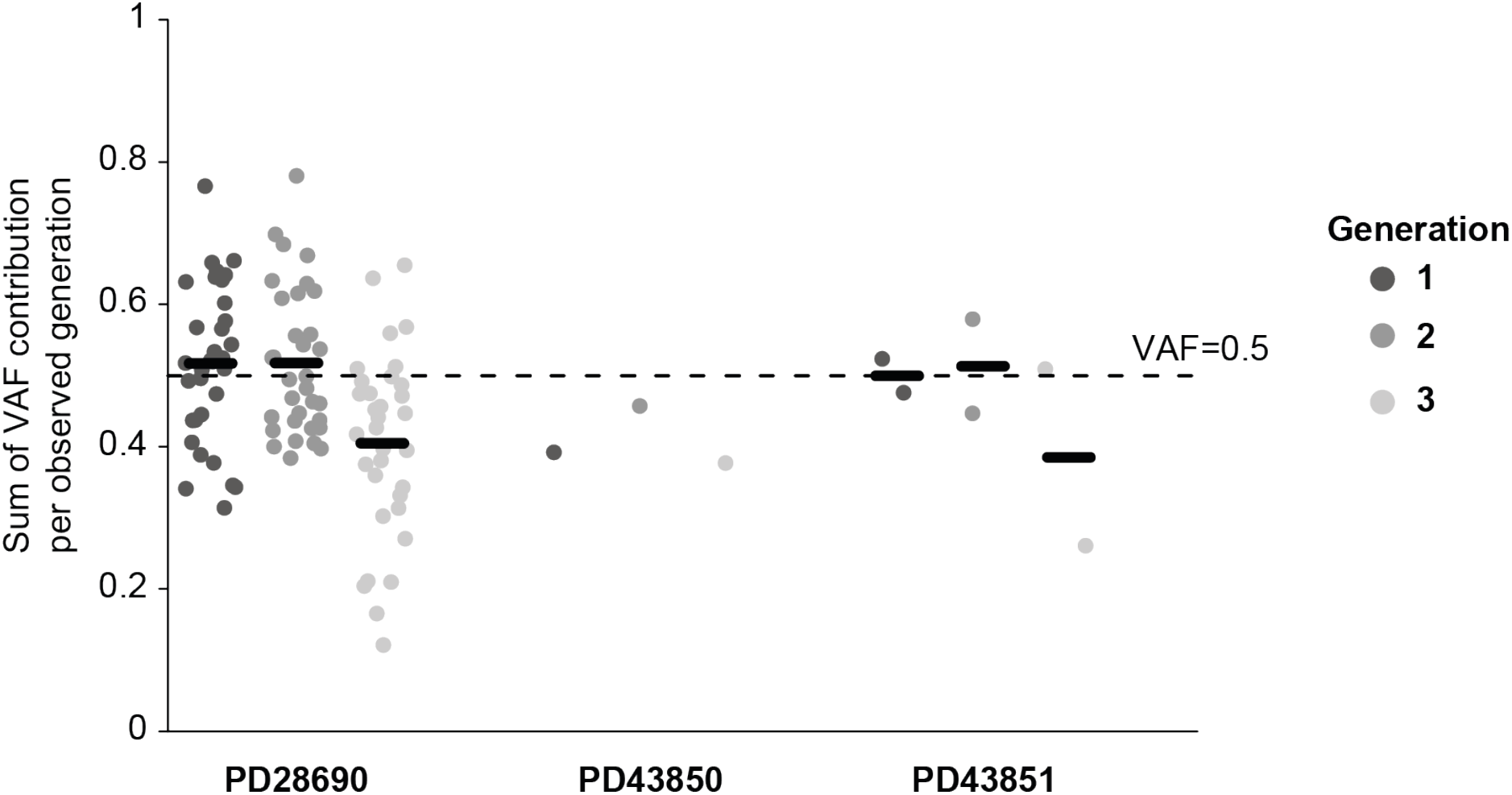
Sum of VAFs per phylogenetic generation. Sum of mean VAFs of branches of the same generation per bulk sample in PD28690 (**a**, n=33), PD43850 (**b**, n=1) and PD43851 (**c**, n=2). A total sum of mean VAFs approximating 0.5 indicates all cells belong to one of the lineages of that generation and are accounted for i.e. no lineages are missing from the phylogeny. This is mostly the case for generations 1 and 2, but the total VAf of generation 3 indicates missing lineages.

**Extended Data Figure 4.**
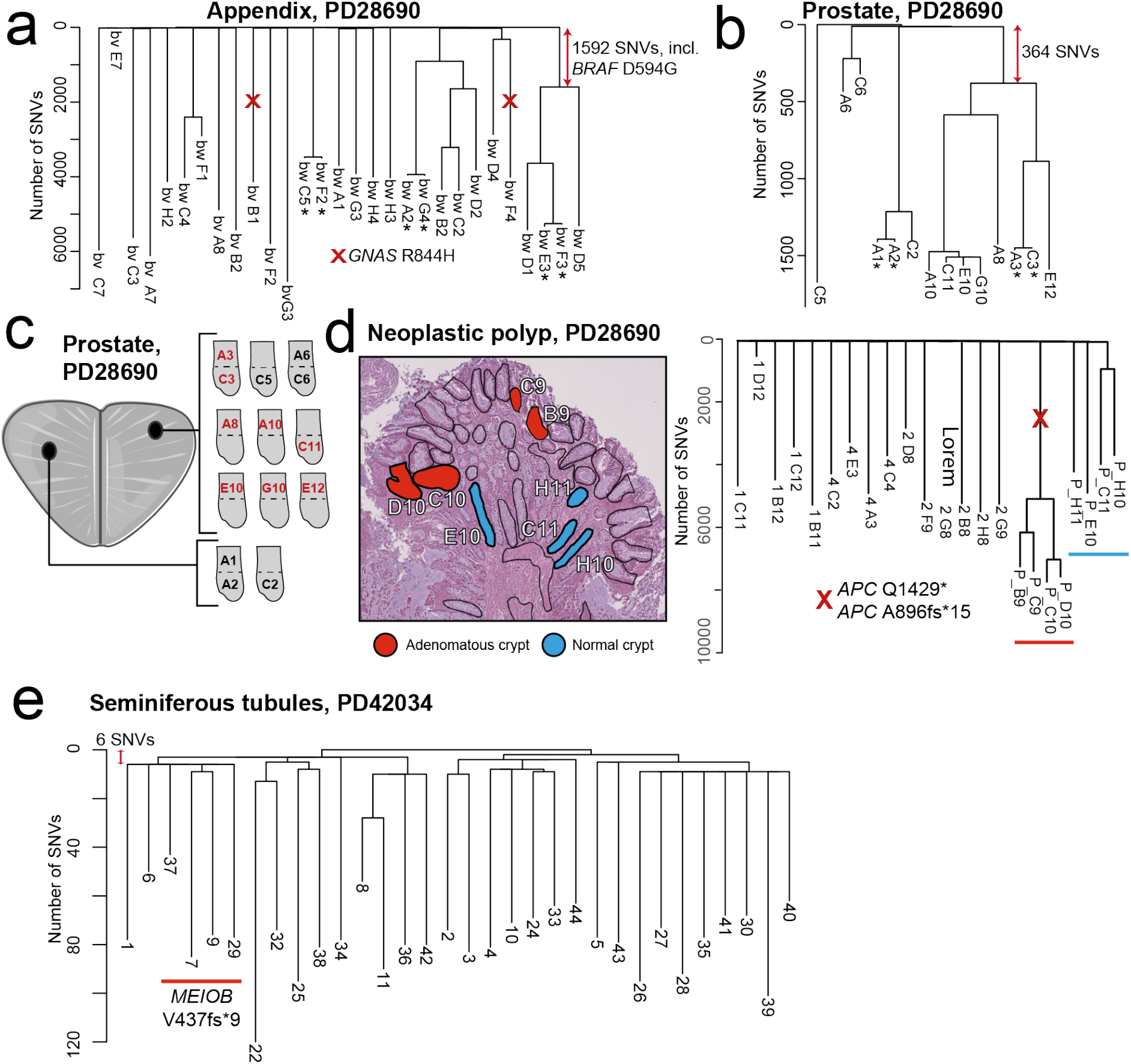
Clonal expansions later in life. **(a**) Phylogenetic tree for appendiceal crypts in PD28690, with annotated cancer driver mutations. An asterisk indicates the two neighbouring crypts were taken as biological replicates of one another. Assuming SNVs have occurred in a linear fashion, the *BRAF* mutation was acquired prior to the age of approximately 23. Phylogeny (**b**) and sampling overview (**c**) for prostatic acini in PD28690, showing widespread benign prostatic hyperplasia in one biopsy. The earliest post-developmental bifurcation in prostate can be timed to an age of approximately 19 years. (**d)**Histology and sampling overview alongside the phylogeny for a microscopic polyp in the colon of PD28690. (**e)** Phylogeny of seminiferous tubules of PD42034, where a frameshift deletion in *MEIOB* was acquired after only 6 post-zygotic SNVs. Parts of the figure are composed of pictures from Servier Medical Art. Servier Medical Art by Servier is licensed under a Creative Commons Attribution 3.0 Unported License (https://creativecommons.org/licenses/by/3.0/).

**Extended Data Figure 5.**
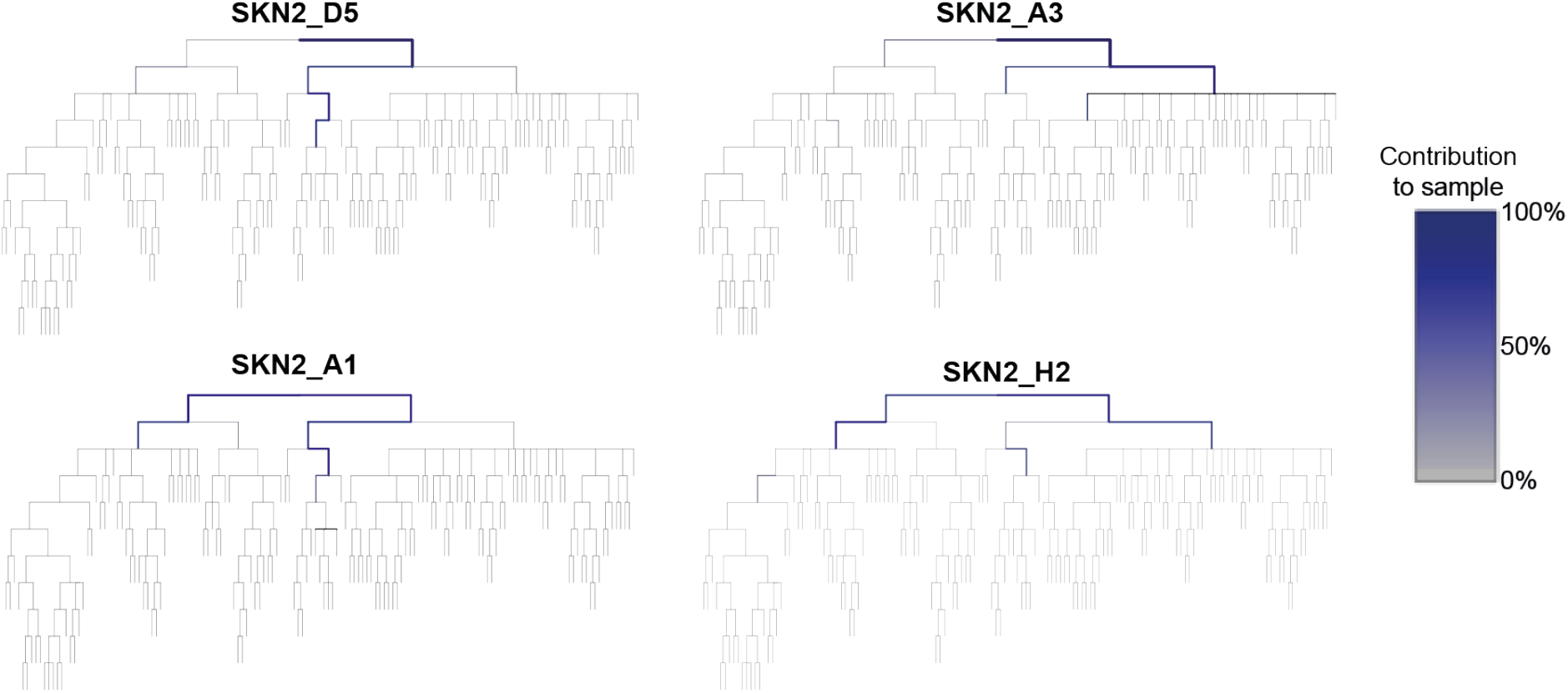
Decomposition of polyclonal samples. Phylogenetic trees with unit branch lengths for four polyclonal samples of epidermis of PD28690, showing the contribution (blue) of early embryonic progenitors in the phylogeny to the sample.

**Extended Data Figure 6.**
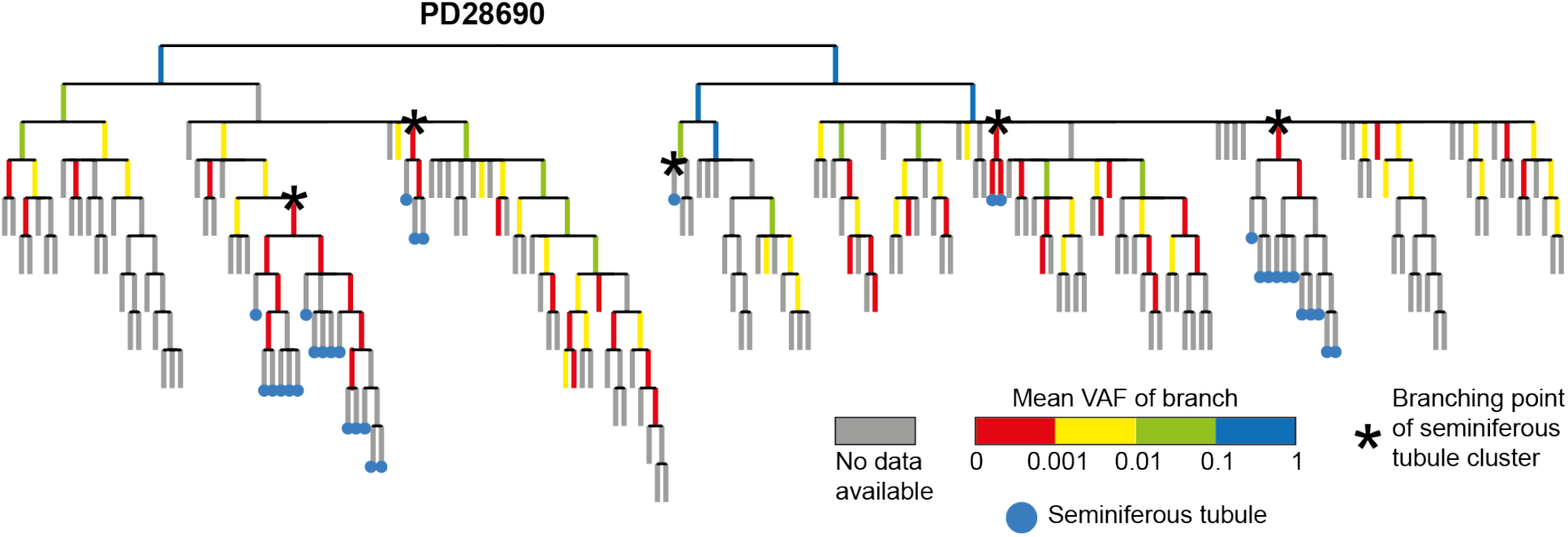
Targeted re-sequencing in PD28690. Cladogram of PD28690 with contribution to 84 bulk samples (none derived from testes) as assessed through targeted re-sequencing of embryonic and spermatogonia-specific variants. The colour of the branch indicates the mean VAF of substitutions on that branch across all bulk samples. Nodes that gave rise to only seminiferous tubules are annotated with an asterisk. Branches coming from those nodes do not contribute to the bulk samples, confirming the segregation of primordial germ cells at the estimated time.

**Extended Data Figure 7.**
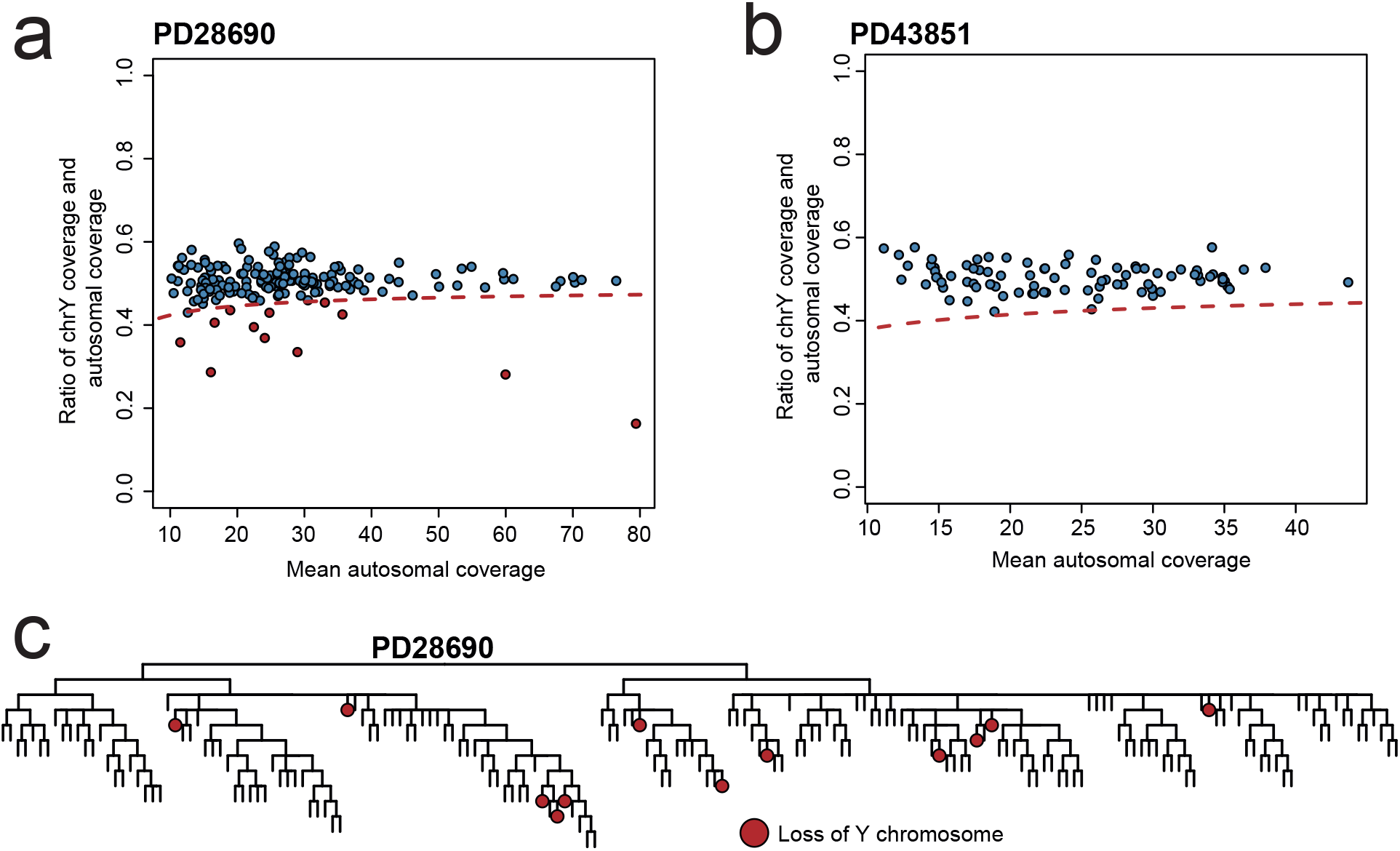
Loss of chromosome Y. Scatterplot showing the ratio between the mean Y-chromosomal coverage and autosomal coverage against the mean autosomal coverage for all samples of PD28690 (**a**) and PD43851 (**b**). The dashed red lines indicates the 95% confidence interval around an expected ratio of 0.5. Red-coloured dots indicate samples with significant evidence of loss of the Y chromosome. (**c**) Phylogeny of PD28690 with samples exhibiting LOY marked in red, indicating all LOY events are acquired independently.

